## Supplemental tables and figures for "OpDetect: A convolutional and recurrent neural network classifier for precise and sensitive operon detection from RNA-seq data"

### OpDetect's Supplementary Material

Rezvan Karaji and
Lourdes Peña-Castillo
Memorial University of Newfoundland

April 7, 2025

#### Contents

**1 Supplementary Tables** **2**

**2 Supplementary Figures** **4**

### 1 Supplementary Tables

Supplementary table 1: List of Python packages with their versions.

| **Package** | **Version** | **Package** | **Version** | **Package** | **Version** |
|---|---|---|---|---|---|
| numpy | 1.24.4 | pandas | 2.1.0 | matplotlib | 3.8.4 |
| scipy | 1.10.1 | tensorflow | 2.16.1 | scikit_learn | 1.2.1 |

Supplementary table 2. Machine learning summary table as per DOME recommendations (https://doi.org/10.1038/s41592-021-01205-4)

| DOME | Version | 1.0 |
|---|---|---|
| **Data** | Provenance | See Table 1 in manuscript. Some data was previously used elsewhere. |
| | Dataset splits | 10-fold cross-validation used to find the optimal hyper-parameters for the CNN-LSTM architecture.<br>See Table 2 in manuscript for number of instances per label in training data. Instances per label in validation set described in Table 6 in manuscript. |
| | Redundancy between data splits | There is not intersection between organisms used in training and validation. To make the split more strict, validation data contains data from a phylum (Spirochaetota*)* not included in the training data, and for a eukaryote (all training data is from bacterial species). |
| | Data availability | Yes. https://github.com/BioinformaticsLabAtMUN/OpDetect |
| **Optimization** | Algorithms | CNN-LSTM architecture. Proposed before by Singh et al. (see complete reference in manuscript) |
| | Meta-predictions | No |
| | Data encoding | RNA-seq read counts across nucleotide bases in a genome. See section *Feature Representation* in manuscript for more details. |
| | Parameters | See Tables 3 and 4 in manuscript. Optimized using 10-fold cross-validation |
| | Features | No feature selection was performed. |
| | Fitting | 10-fold cross-validation was used to find the optimal hyper-parameters for the CNN-LSTM model.<br>Independent validation data used to further evaluate performance. |
| | Regularization | Dropout and early stopping. |
| | Model availability | Yes. https://github.com/BioinformaticsLabAtMUN/OpDetect |
| **Model** | Interpretability | Black box. |
| | Output | Classification – probability of pair of genes belonging to the same operon. |
| | Execution time | In a high-performance computing environment, executing times are as followed:<br>• 10-fold CV (training) 1h23m, RAM 1.7 GB, 4 cores<br>• Prediction for ~5.6k gene-pairs requires ~3m20s, RAM 540MB and 1 core. |
| | Availability of Software | Yes. https://github.com/BioinformaticsLabAtMUN/OpDetect |
| **Evaluation** | Evaluation method | Cross-validation.<br>Independent dataset. |
| | Performance measures | F1-score, AUROC, recall (see Tables 7 and 8, Figs. 1-3, Supplementary figures 1-7). |
| | Comparison | OperonSEQer, Operon Finder, Operon-mapper and Rockhopper (see Section *Comparative assessment* in manuscript).<br>Methods selected are the most recent machine learning-based ones. |
| | Confidence | Confidence intervals were calculated for the cross-validation results (Table 5). Performed Friedman test, and all-pairs comparisons of AUROC using several pairwise post hoc tests (i.e., Miller, Nemenyi, Siegel and Quade). |
| | Evaluation availability | Code to perform evaluation available at https://github.com/BioinformaticsLabAtMUN/OpDetect |

### 2 Supplementary Figures

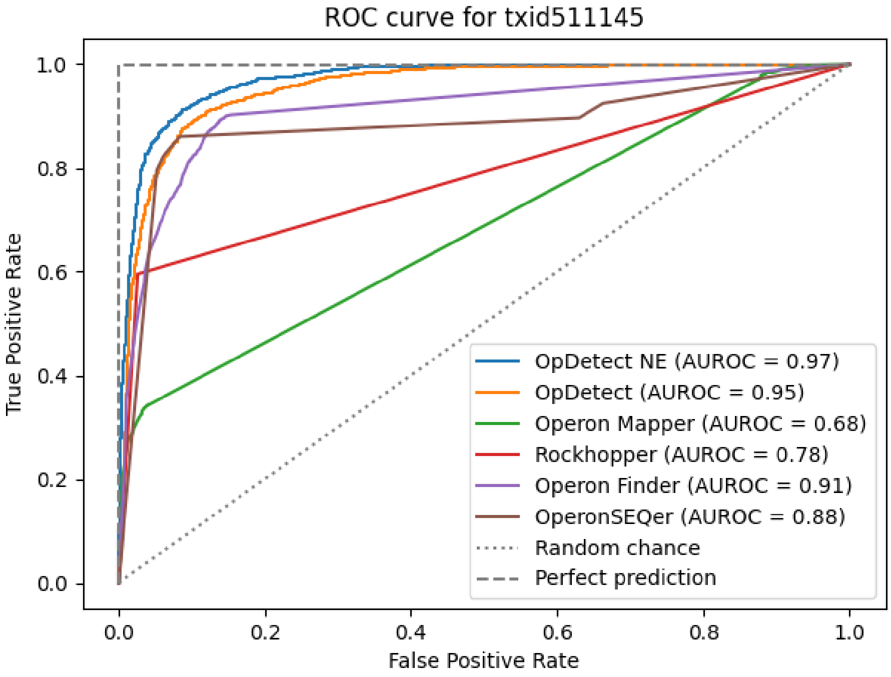


Supplementary figure 1: ROC for *E. coli* K-12 substr. MG1655. "OpDetect NE" refers to OpDetect without excluding the examined organism from the training process (i.e., affected by data leakage).

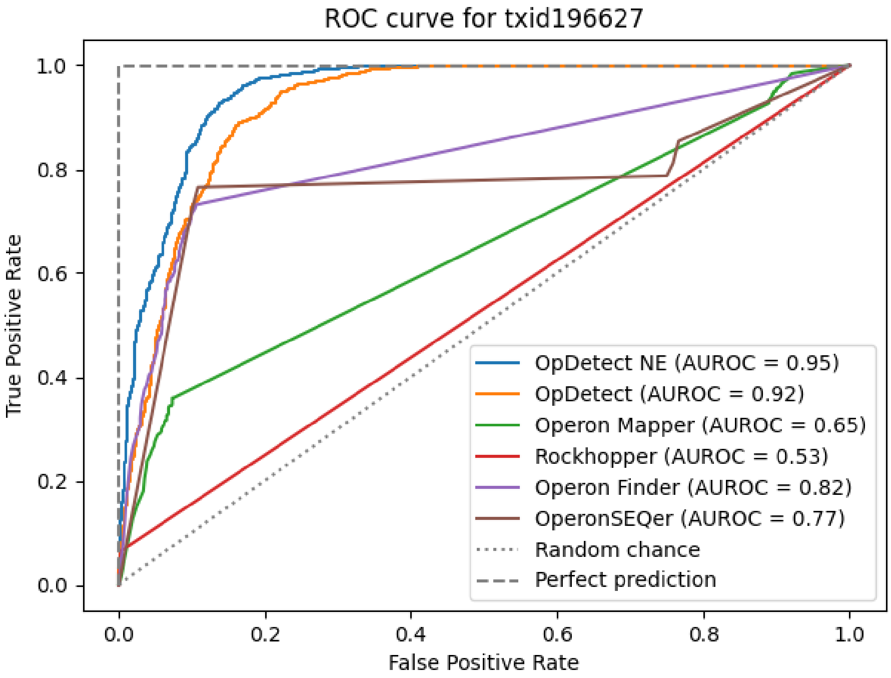


Supplementary figure 2: ROC for *C. glutamicum* ATCC 13032. "OpDetect NE" refers to OpDetect without excluding the examined organism from the training process (i.e., affected by data leakage).

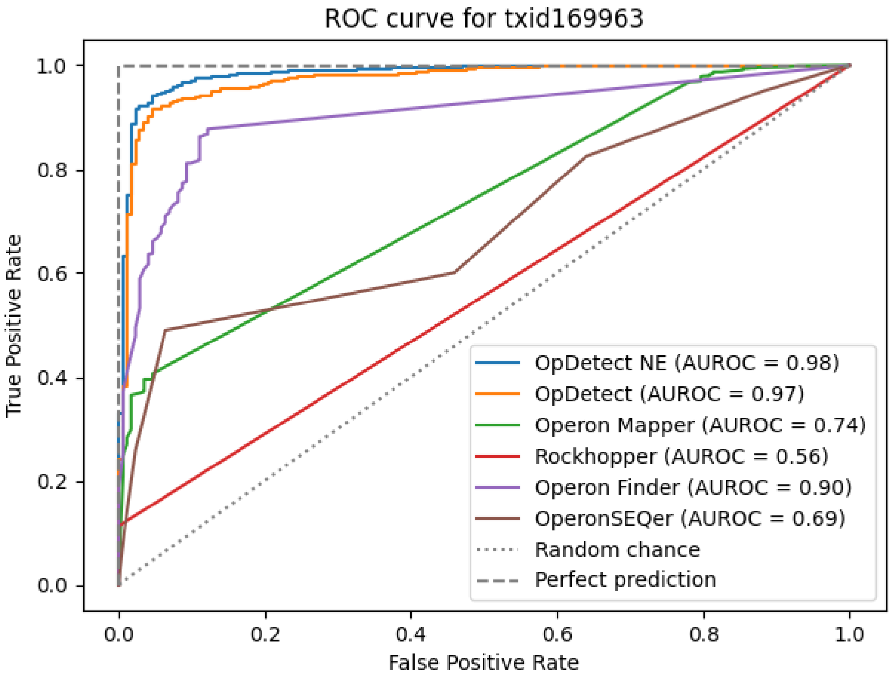


Supplementary figure 3: ROC for *L. monocytogenes* EDG-e. “OpDetect NE” refers to OpDetect without excluding the examined organism from the training process (i.e., affected by data leakage).

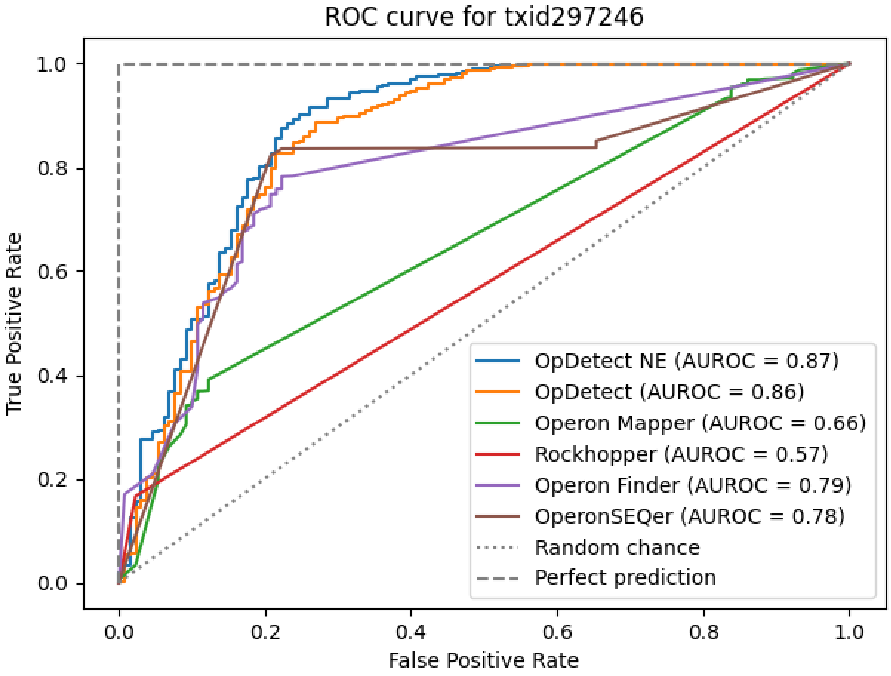


Supplementary figure 4: ROC for *L. pneumophila* str. Paris. "OpDetect NE" refers to OpDetect without excluding the examined organism from the training process (i.e., affected by data leakage).

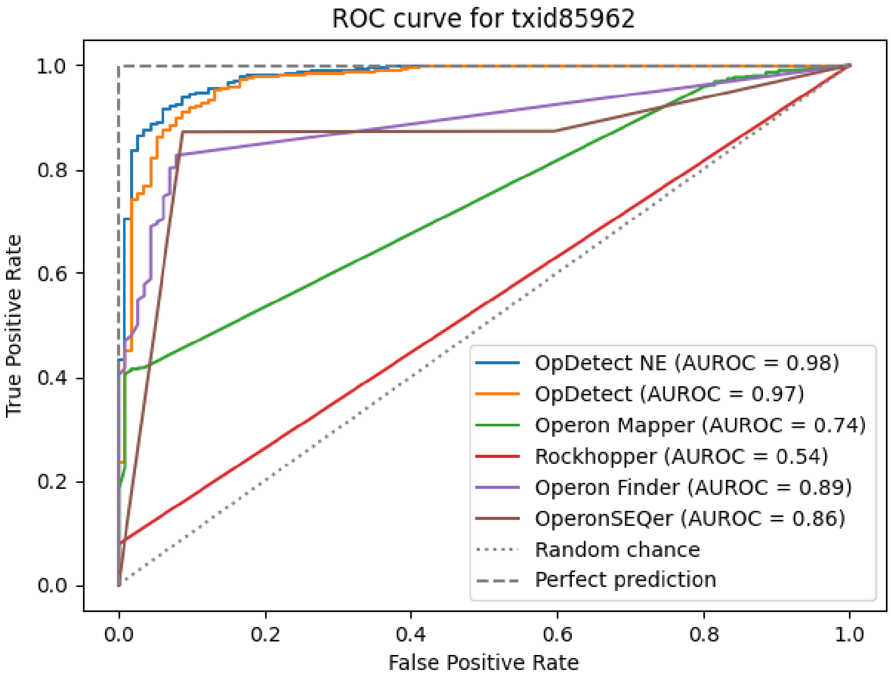


Supplementary figure 5: ROC for *H. pylori* 26695. "OpDetect NE" refers to OpDetect without excluding the examined organism from the training process (i.e., affected by data leakage).

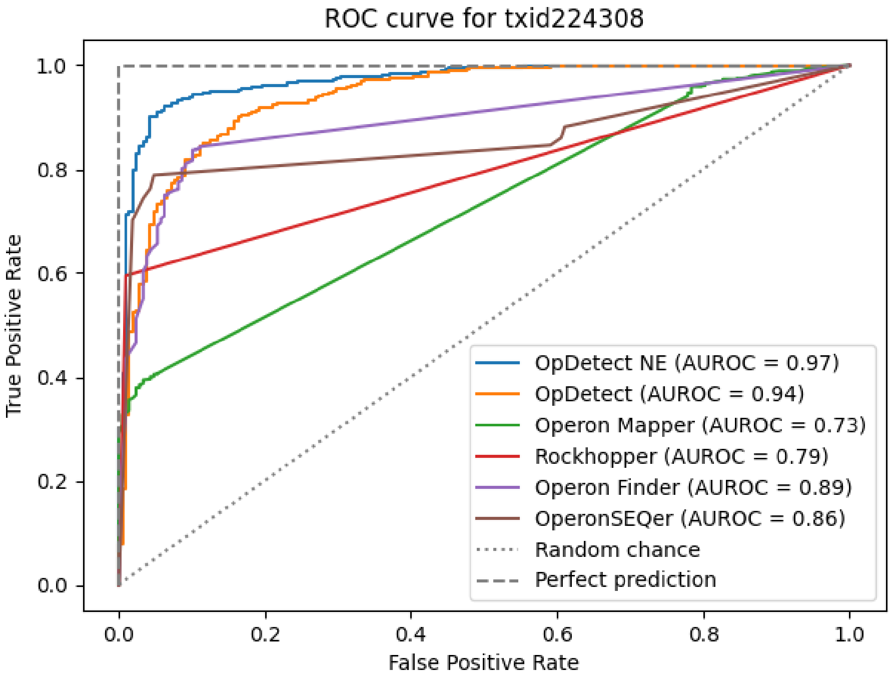


Supplementary figure 6: ROC for *B. subtilis* subsp. subtilis str. 168. "OpDetect NE" refers to OpDetect without excluding the examined organism from the training process (i.e., affected by data leakage).

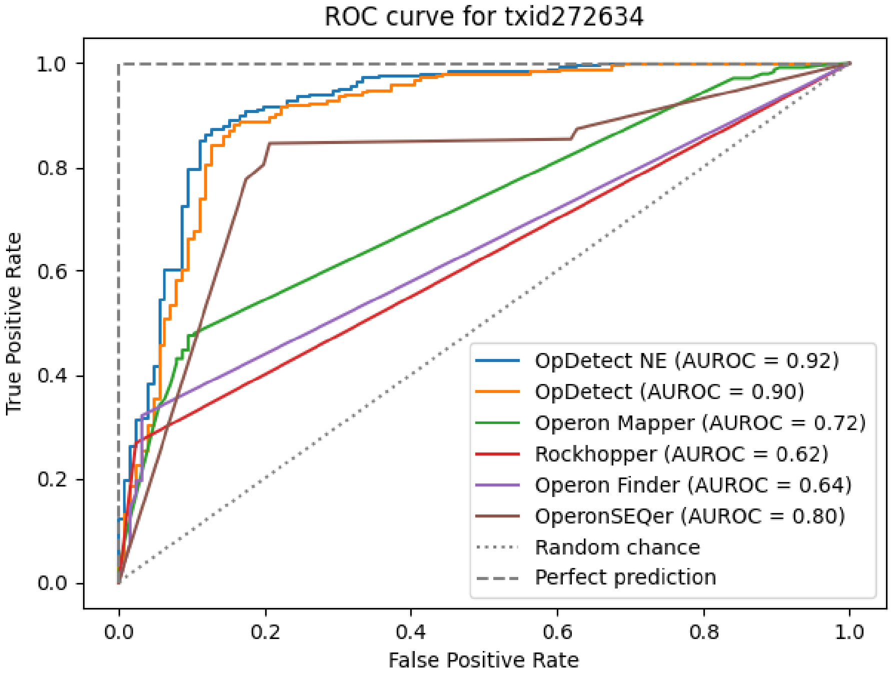


Supplementary figure 7: ROC for *M. pneumoniae* M129. "OpDetect NE" refers to OpDetect without excluding the examined organism from the training process (i.e., affected by data leakage).
